# Machine learning-assisted directed evolution yields dramatic improvement on novel AAV engineering task

**DOI:** 10.64898/2026.10.06.757046

**Authors:** Marina Luchner, Ioulia Vorobieva, Ruhani Makkar, Mattea Jozinovic, Simon Chester, Matthew JA Wood, Stephan J. Sanders, Dhanu Gupta, Harrison Steel

**Affiliations:** Department of Engineering Science, University of Oxford, Parks Road, Oxford, OX1 3PJ; Department of Paediatrics, University of Oxford, Children’s Hospital, John Radcliffe, Headington, Oxford, OX3 9DU, UK; Institute of Developmental and Regenerative Medicine, University of Oxford, IMS-Tetsuya Nakamura Building, Old Road Campus, Roosevelt Dr, Headington, Oxford OX3 7TY, UK; Department of Bioscience, TUM School of Natural Sciences, Technical University of Munich, Boltzmannstr. 10, 85748 Garching; Department of Psychiatry and Behavioral Sciences, UCSF Weill Institute for Neurosciences, University of California, San Francisco, San Francisco, CA 94158, USA2,3; Biomolecular Medicine, Division of Biomolecular and Cellular Medicine, Department of Laboratory Medicine, Karolinska Institutet, Huddinge 14151, Sweden

## Abstract

Directed evolution enables the discovery of protein mutants with improved fitness through iterative rounds of selection and has been widely applied to adeno-associated virus (AAV) capsid engineering. Machine learning (ML) can augment this process by prioritising mutants for experimental validation, but whether its benefits outweigh the added cost of ML-designed library construction remains unclear. Here, we address this question by considering improvements in manufacturing efficiency of AAV capsids in the context of exosomal encapsulation, which is a technique for improving AAV immunogenicity. We generated a directed evolution dataset comprising 53,974 mutants across three rounds of selection and an independent assessment dataset of 472 ML-designed mutants with detailed profiling. Directed evolution alone yielded a 3-fold improvement, while ML-assisted directed evolution yielded a 41-fold improvement in manufacturing efficiency, demonstrating that ML-assisted directed evolution outperforms directed evolution alone. We show that data quality – specifically the number of selection rounds and the statistical power of mutant counts – has a greater impact on model performance than model architecture. Finally, we provide practical guidance on experimental design and implementation of ML tuning strategies that can augment model performance in protein engineering applications.

## Introduction

Protein engineering is essential for the effective manufacturing of vehicles for gene therapy delivery (including viral vectors, virus-like particles and exosomes)^1^. However, the design space for protein sequences is typically too large to search exhaustively in the laboratory. This motivates the application of directed evolution, in which successive rounds of mutagenesis, measurement, and selection can identify mutants with improved characteristics^2^. The effectiveness of directed evolution can be augmented through machine learning, steering the search through protein design space via the addition of computational *learning* and *prediction* steps^3,4^. Such approaches are especially promising for exploring protein fitness landscapes that are challenging for traditional directed evolution^5^. While ML-assisted methods are thus promising in principle, significant open questions remain regarding the quantitative benefit they may achieve in practice – and how this might be maximised.

To address this challenge, we benchmark ML-assisted directed evolution against standard directed evolution on a challenging and novel adeno-associated virus (AAV) engineering task, which aims to address key limitations for gene therapy delivery. AAVs are a standard delivery vehicle for gene therapies, with 7 AAV-based gene therapies licensed to date^6^. Directed evolution has been applied to engineer AAV capsid protein for improved titres and immune evasion^7,8^, two factors amongst others that are currently limiting clinical translation. To evolve AAVs, large libraries of capsid mutants are created and selected to enrich those with desired phenotypes^9,10^. A notable success of directed evolution was AAV-PHP.B, an AAV9 mutant that can cross the blood–brain barrier of adult mice for widespread central nervous system transduction^11^. While directed evolution has been successfully applied to improve AAV for gene therapy purposes, the throughput of experimental methods leaves a large fraction of the protein design space unexplored (e.g., an 8-mer insertion corresponds to ∼25.6 billion possible mutants), and maximisation of sequence fitness can be locked in local rather than global maxima.

Here, ML offers a promising path forward^12^: Supervised models can be trained on high-throughput screening data to predict AAV capsid performance directly from sequence, enabling in silico screening of new mutants^13^. More recently, ML has been leveraged to advance AAV engineering beyond traditional approaches. ML-guided library design for AAV peptide insertions achieved fivefold higher virus titres than a standard NNK random library^14^, accurately predicted capsid functionality in macaque monkeys from *in-vitro* data^8^, and yielded mutants with a greater average diversity than natural AAV serotypes^15^. Model architectures tested on AAV data include neural networks^8,14,15^, protein language models^16^ and variational autoencoders^17^, underscoring the broad interest of ML-assisted design within AAV engineering. AAV encapsulation into exosomes (EVs), is a novel engineering strategy that has recently been shown to address immunogenicity and transduction efficiency concerns^18^. EV-AAVs have been shown to outperform AAVs in delivering a genetic payload to the murine retina^19^ and mediating delivery to the CNS after systemic injection in mice, even for serotypes that typically do not cross the blood-brain barrier^20^.

While outperforming AAVs in this regard, EV-AAVs present additional experimental challenges that must be overcome to allow their widespread application. Because only a fraction of AAVs are encapsulated within EVs during the manufacturing process, efficient scale-up and translation of this vector to clinical applications has proven challenging. Previous work attempted to overcome low yields by overexpressing CD9, an EV biomarker, and MAAP, an AAV protein involved in cellular membrane association. However, these approaches only achieved modest 2- to 3-fold improvements in manufacturing efficiency^21,22^. Other methods aim to improve efficiency through cellular nanoporation and metabolic engineering, which increases the quantity of EVs provided for encapsulation *rather than* optimising the percentage of AAVs that are encapsulated^23,24^.

Here, we developed a novel ML-assisted directed evolution approach to evolve AAV capsid mutants with enhanced EV encapsulation efficiency, employing a range of ML models (from linear regression to deep learning architectures), and benchmarking this approach against a standard (non-ML) assay. We evolved AAV capsid mutants with a 40.9-fold improvement in overall EV-AAV manufacturing efficiency and encapsulation efficiencies of up to 99.2% – a result on the scale necessary to close the gap in manufacturing capability between AAV and EV-AAV vectors^21^. Beyond implications for the field of gene therapy, our results reveal the magnitude of improvement that ML-assisted directed evolution can deliver compared to traditional directed evolution.

## Results

### Scalable characterisation of AAV capsid mutants

Our experimental approach implements and compares two directed evolution methods; a “traditional” directed evolution assay (Fig.1a, top), and an ML-assisted directed evolution approach in which experimental observations are used to design and test novel mutants that (may) perform better than those observed initially (Fig. 1a, bottom). In both cases, the engineering pipeline is initiated with a mutant library, created by inserting eight to thirteen random amino acids (8-mer, 9-mer, 12-mer, 13-mer) between residues 453-454 into a surface-exposed loop of the AAV9 capsid protein^25^, leveraging sites that tolerate insertions without compromising assembly or infectivity. Random NNK sequences (where N represents any nucleotide and K represents either G or T) were cloned into the AAV genome backbone, generating pre-encapsulation libraries with a theoretical minimum complexity of ∼25.6 billion mutants at the amino acid level (20^8^ amino acid combinations for the 8-mer amino acid inserts). Note that building sequences with NNK codons is a common strategy to reduce the frequency of non-functional sequences that contain stop codons^26–28^.

**Figure 1:**
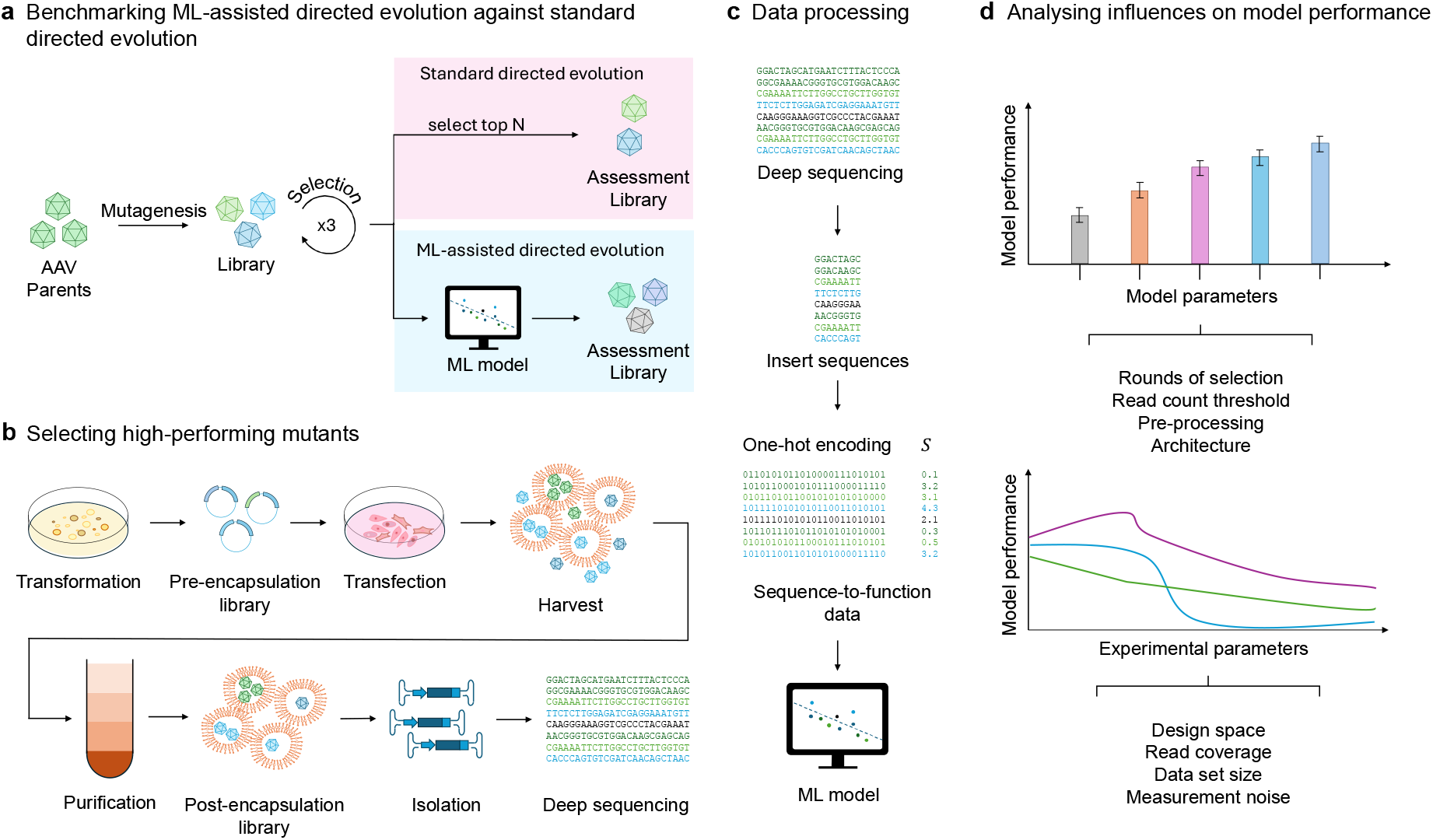
ML-assisted directed evolution pipeline and optimisation demonstrated on a novel protein engineering task. (**a**) “Standard” directed evolution of AAV capsid protein involves mutagenesis and three cycles of selection. ML-assisted directed evolution uses data generated during different rounds of selection to train ML models, which are used to predict protein mutants with enhanced properties. Here, we are benchmarking ML-assisted directed evolution against a standard directed evolution method, which selects the top *N* observed mutants. (**b**) Selecting for enhanced EV encapsulation properties involves cloning the gene encoding AAV capsid protein into *Escherichia coli* on a plasmid, isolating and transfecting the plasmid into HEK 293 T cells and harvesting the supernatant. Next, the EV-AAVs are isolated using iodixanol gradient purification. After isolating the AAV genome from within the EV-AAVs, samples are sent for deep sequencing. (**c**) Dataset generation requires processing the raw sequencing data, deriving the read count per mutant, and calculating the log-transformed enrichment score *S* of the mutant in the post-encapsulation compared to the pre-encapsulation library, yielding a sequence-to-function dataset. The ML model is then trained to predict the log-transformed enrichment score *S* from the one-hot encoded insert sequence. (**d**) We analysed the performance of different tuning strategies as well as the influence of different experimental parameters on ML model performance.

This library then underwent three rounds of selection to enrich high-performing mutants by encapsulating AAVs into EVs, harvesting and purifying EV-AAVs, and re-cloning selected mutants to amplify them for subsequent rounds (Fig.1b). Following each selection step, purified EV-AAVs were treated with DNase and Protease K to degrade surface-bound DNA and isolate encapsulated genomes, yielding three pre- and post-encapsulation libraries (Methods). Finally, deep sequencing enabled quantitative assessment of the EV-AAV manufacturing efficiency (i.e. our protein fitness metric-of-interest) defined in terms of the enrichment score *S* of the AAV genomes in the post-encapsulation compared to the pre-encapsulation library^22^ (Fig.1c). For the first and third round of selection, we measured enrichment scores *S* for 54K and 18K AAV capsid mutants, respectively. However, these datasets only cover a small fraction of the theoretical complexity, motivating the need for exploring the remainder of the protein design space.

To compare ML-assisted directed evolution against “traditional” directed evolution, our workflow diverged at this point. For “traditional” directed evolution, the best 123 mutants observed in the enrichment library were selected, re-synthesised, and measured in a confirmatory assessment experiment. In contrast, the ML-assisted directed evolution approach used the data generated throughout the three rounds of selection to train an ML model, which was then used to generate an assessment library of 472 mutants with strong predicted fitness.

### Model training and evaluating predictive performance

To create ML models that predict each mutant’s EV-AAV manufacturing efficiency based on amino acid sequence, we adapted the framework developed by Yang et al.^3^: following each round of directed evolution and (with the first and third round referred to as Round_1 and Round_3), deep sequencing of enriched mutants is used to measure EV-AAV manufacturing efficiency data sets for model building (training, tuning hyperparameters, testing). In this model, the one-hot encoded amino acid sequence acts as model input, and the log-transformed enrichment score *S* (assessing the EV-AAV manufacturing efficiency) represents the model output (see Methods). Model performance is quantified by calculating the Pearson’s correlation between ML-predicted and measured log-transformed enrichment score *S* on a held-out test set (Fig. 2, Supplementary Fig. S1). In total, we trained and compared the performance of fifteen classes of ML models: six linear regression (LR) models, five long short-term memory (LSTM) models, two feed-forward neural network (FNN) models, and two convolutional neural network (CNN) models (Supplementary Table S1).

**Fig. 2:**
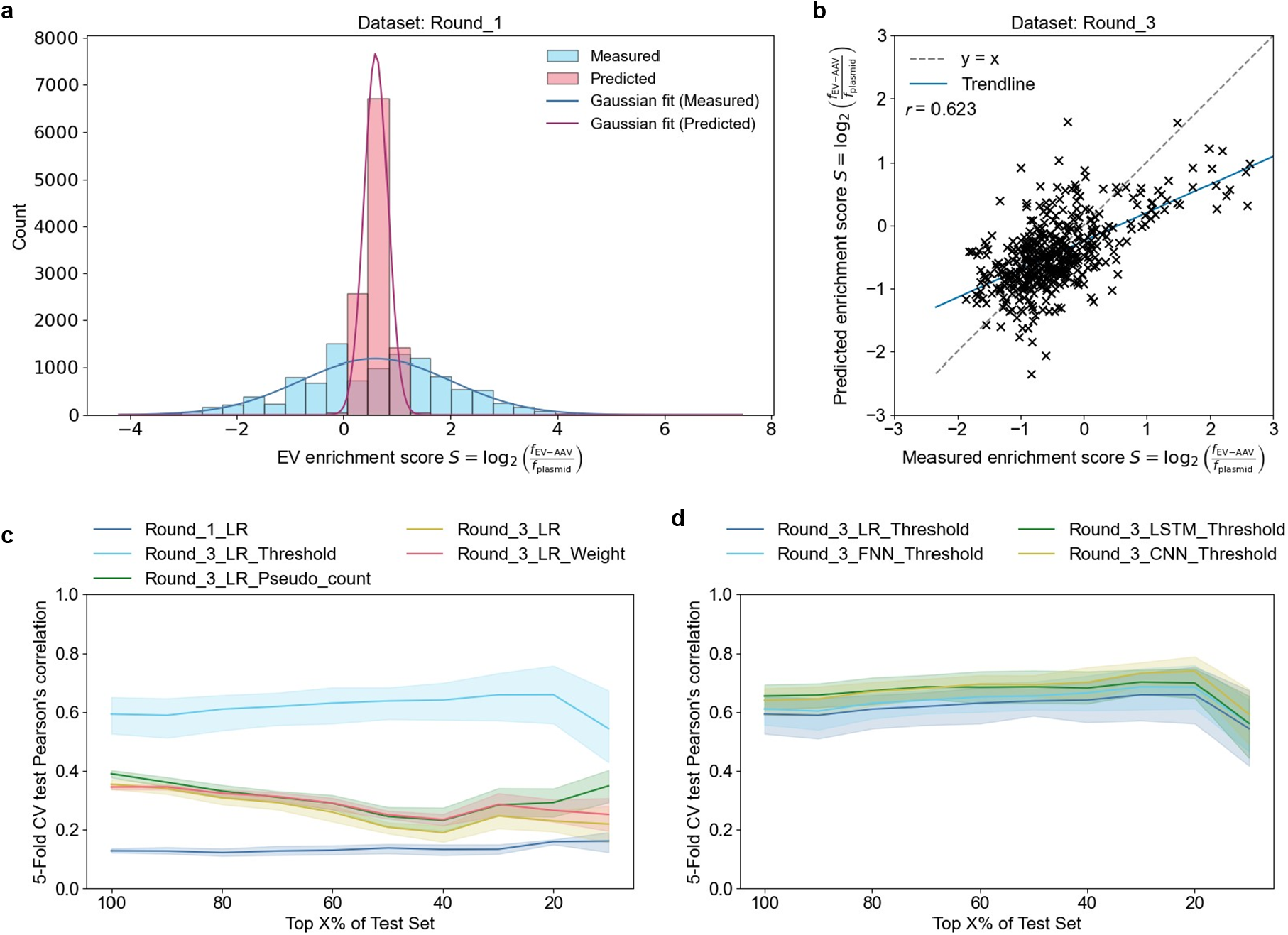
Tuning ML model performance to learn EV recruitment from amino acid sequence. (**a**) Overlap of predicted and measured (“true”) EV enrichment score *S* distributions from data generated in the first round of selection (Round_1). (**b**) Correlation between model predictions and experimentally measured EV enrichment scores *S* from data derived during the third round of selection (Round_3), excluding *S* by thresholding the read count. (**c**) Sensitivity analysis of model performance when ablating the test data to retain top N% of EV enrichment scores *S* plotted with 95% confidence intervals. Pre-processing strategies performed on Round_3 include accounting for statistical power of the EV enrichment score *S* (Round_3_LR_Weight), filtering high quality scores (Round_3_LR_Threshold) and imputing missing values (Round_3_LR_Pseudo_count). (**d**) Sensitivity analysis of model performance using Round_3 and applying thresholding. Models include linear regression (Round_3_LR_Threshold), a long short-term memory model (Round_3_LSTM_Threshold), a convolutional neural network (Round_3_CNN_Threshold) and a feed-forward neural network (Round_3_FNN_Threshold). Average Pearson’s correlation was calculated on a held-out test set under five-fold cross-validation (CV) and plotted ±95% confidence intervals.

First, we compared the influence of different pre-processing strategies on the performance of our baseline LR model; these strategies included weighting *S* based on its statistical power (Round_3_LR_Weight), excluding *S* by thresholding the read count (Round_3_LR_Threshold), and replacing missing read counts with a pseudo count of 1 (Round_3_LR_Pseudo_count). Each pre-processing strategy improved model performance, with thresholding yielding the greatest improvement (Fig. 2c), despite reducing the training dataset size from 16K to 2K.

As our primary goal was the selection of *top*-performing mutants, we conducted a sensitivity analysis in which the randomly held-out test set was restricted to its top N% of mutants by measured enrichment score. We found that model performance was maintained, and in some cases even improved, when only the top 20% of mutants were included in the test set (corresponding to between 1K and 40 mutant sequences depending on the pre-processing strategy). We observed that using data from the final round of selection and conditioning inclusion of training data by read count thresholding led to markedly improved model performance (see Round_3_LR_Threshold in Fig. 2c), resulting in a Pearson’s correlation coefficient of 0.62 under five-fold cross-validation.

Next, we investigated the effect of differing model architectures on model performance, including LR and models allowing for more complex relationships, such as LSTMs and CNNs. CNNs, for example, have previously demonstrated strong performance in predicting protein expression from sequence, specifically in regimes characterised by sparse data^29^, and convolutions have been shown to be competitive with transformers in the protein prediction context^30^. Across all architectures, model performance notably plateaued at a Pearson’s correlation coefficient of approximately 0.65 when tested on the complete held-out test set. This Pearson’s correlation represents an increase compared to previous methods that report correlations of up to 0.59 on a related AAV capsid engineering task^10^. Interestingly, different model architectures show similar performance within the bounds of 5-fold cross-validation (Fig. 2d). Training and validation loss curves show the same pattern: all architectures converged within 50 epochs to comparable validation loss (Supplementary Fig. S2a), consistent with architecture being a weak determinant of performance in this regime.

To further analyse whether hyperparameter tuning has an impact on model performance, we tested varying numbers of hidden layers and nodes for the LSTM architecture, which has been chosen as the preferred model for a similar AAV engineering task^8^. Surprisingly, variations in model depth and the number of nodes per hidden layer had no statistically significant effect on predictive performance (Supplementary Fig. S3a). Together, these results suggest that parameters other than model architecture and hyperparameters constitute the primary bottleneck for model performance. The significant improvements in model performance through thresholding low-quality reads and multiple rounds of selection suggest that experimental parameters, such as measurement noise, are the main limitation. Consequently, we selected Round_3_LR_Threshold for further evaluation, as it performed similarly to the overall best-performing model Round_3_CNN_Threshold, while offering greater interpretability.

To rule out that Round_3_LR_Threshold performance estimates are skewed by data leakage, we repeated model evaluation under a similarity-aware split. Because every 8-mer occurs exactly once in the dataset, the training and test sets share no sequences, and the highest sequence identity between them is 87.5% – a single mismatch across the eight positions – which only 32 of 404 test sequences reach (Supplementary Fig. S4a). Constraining test sequences to fall below successively stricter identity thresholds to all training sequences left predictive performance essentially unchanged (Pearson’s *r* = 0.62, 0.66, 0.52 and 0.59 at cutoffs of 87.5%, 75%, 62.5% and 50%, respectively; Supplementary Fig. S4b), indicating that the reported correlations are not driven by data leakage between training and test datasets.

### ML-assisted directed evolution yields top mutants with markedly higher fitness compared to standard directed evolution

With an effective model selected, we sought to compare the EV-AAV manufacturing efficiency of ML model-designed sequences with that of sequences identified as the most enriched during the third round of selection, providing an estimate for the added value of ML for directed evolution workflows. To form an assessment library for the ML-assisted approach, we generated novel mutants using Round_3_LR_Threshold. The choice of a linear model is more broadly motivated because neural network models did not provide a significant advantage, LR is less prone to overfitting, and sequence design with LR is straightforward and interpretable. Specifically, the LR model assigns a weight *w* to every (insert position *p*, amino acid *a*) pair and adds a bias *b* to predict the EV enrichment score *S* from the input features *x*_*p,a*_ (Equation 1). The input features *x*_*p,a*_ are one-hot encoded, meaning that *x*_*p,a*_ = 1 if amino acid *a* is present at position *p* and otherwise *x*_*p,a*_ = 0.

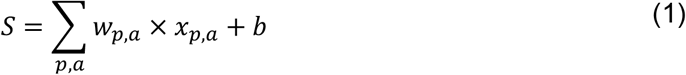

When inspecting *w*_*p,a*_ of Round_3_LR_Threshold, we noticed that cysteine was assigned the largest weights across insert positions two to seven. Therefore, designing novel sequences purely based on the maximum predicted fitness would yield sequences with sequential cysteines. However, sequential cysteines represent a concern for experimental protein engineers due to their elevated chemical reactivity. Furthermore, a library with maximum predicted fitness would be of low sequence diversity, which can be undesirable^14^. To balance sequence diversity and predicted fitness while avoiding sequential cysteines, we used Round_3_LR_Threshold to generate a position-wise probability distribution for high-performing amino acid sequences. We randomly sampled from this distribution rather than selecting maximum-fitness sequences, yielding diverse sequences with high predicted average performance (see Methods). This strategy enables exploration of the local fitness landscape while avoiding premature convergence on a narrow subset of highly similar sequences, a common limitation of purely greedy optimization approaches^31^.

Sequences identified by both ML-assisted and “standard” directed evolution were synthesised, forming assessment libraries, and their enrichment score *S* quantified (as in Fig. 1b). The top sequences in the ML assessment library reached a mean *S* of 83.1 across three replicates, a 15.1-fold increase over the top-ranked position in the directed evolution assessment library (*S* = 5.5; Fig. 3a). This increase was significant under a one-sided Mann-Whitney U test with p ≤0.05. Relative to wild-type AAV9 measured in the same screen (*S* = 2.03), this represents a 41-fold improvement in EV-AAV manufacturing efficiency, compared with the 3-fold improvement that standard directed evolution achieved over wild-type in its own screen. An improvement of this magnitude is comparable to the reported gap in manufacturing capability between EV-AAVs and AAVs^21^.

**Fig. 3:**
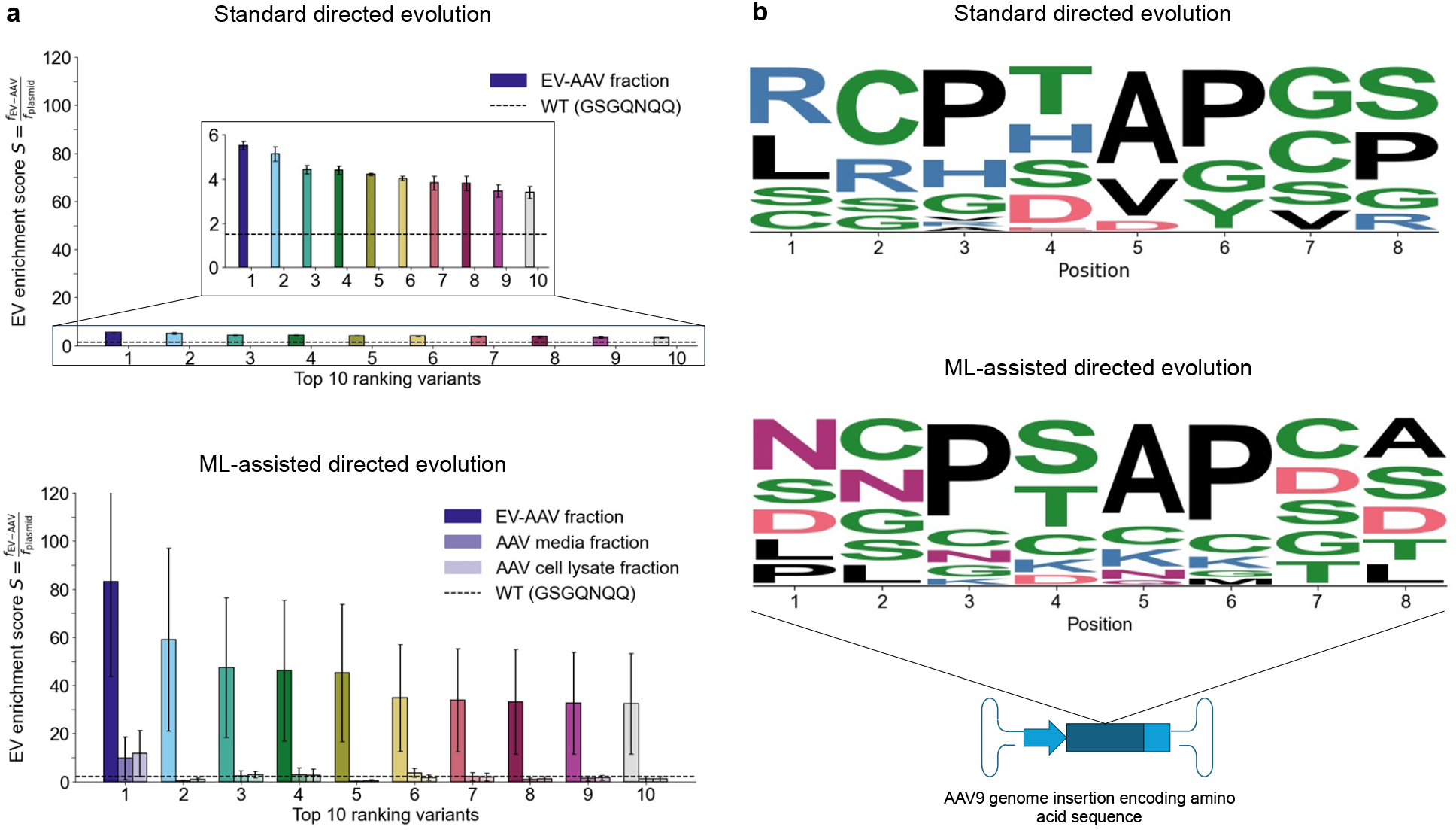
Benchmarking standard directed evolution against ML-assisted directed evolution. (**a**) AAV capsid mutants were selected using either approach, forming assessment libraries. Plasmids encoding AAV capsid mutants were assembled, amplified in Escherichia coli, and pre-encapsulation libraries were transfected into HEK 293 T cells. EV-AAV and AAV particles secreted from the producer cells accumulated in the supernatant, while the remaining AAV particles were retained intracellularly. EV-AAV and AAV fractions were isolated from the supernatant (media), and AAV fractions from the cell lysate. Then, AAV genomes were isolated, forming post-encapsulation libraries. Libraries were sequenced, revealing the AAV capsid mutant counts. Enrichment in the post-encapsulation library compared to the pre-encapsulation library was measured (and quantified as enrichment score *S*). For data analysis, AAV capsid mutants were ranked based on *S*, processing each replicate individually, and the average enrichment score *S* for the top 10 ranking mutants was plotted ± standard error. (**b**) Amino acid probabilities per insert position were derived and plotted based on the relative frequency observed across the assessment libraries.

We further evaluated the robustness of the fitness improvement conferred by ML assistance while accounting for differences in assessment library size. Specifically, we found that the top-ranking mutants in each replicate of the ML assessment library consistently outperformed the top mutants in each replicate of the directed evolution assessment library, even when the ML assessment library was subsampled to match the number of mutants as in the directed evolution assessment library (Supplementary Fig. S5). Moreover, the increase in *S* remained significant under a one-sided Mann-Whitney U test with p≤0.01. Since the ML-designed library outperforms the directed evolution assessment library at equivalent size, in future research efforts, the ML-designed library could be screened *instead* of the top mutants selected by directed evolution, requiring no additional experimental resources. ML assistance is not cost-free in absolute terms – the models depend on the three rounds of selection and deep sequencing described above – but its advantage lies in extracting further value from data that a standard directed evolution campaign already produces.

To analyse whether improved EV-AAV manufacturing efficiency stemmed from increasing total AAV titres, we quantified AAV genome enrichment in the supernatant (secreted AAVs) and the lysed producer cells (intracellular AAVs) for the ML assessment library. The top-ranking mutants exhibited higher total AAV genome titres compared to lower-ranking candidates. They were also more efficiently encapsulated: across the ten top-ranked positions, 75.0 - 99.2% of AAV genomes were recovered in the EV-AAV fraction (mean 85.8%), compared with 29.6% for wild-type AAV9 (Supplementary Fig. S6a). Plotting the EV enrichment score *S* against the percentage of encapsulated genomes shows that the two properties vary independently and facilitates the selection of variants with an optimal trade-off (Supplementary Fig. S6b).

We further compared the amino acid sequence profiles between our two directed evolution approaches (Fig. 3b). Notably, the PS/TAP motif, which has been implicated in EV recruitment previously^32^, is strongly represented in both groups. Meanwhile, cysteines are enriched in positions 2 to 7 in the ML assessment library and in positions 2 and 7 in the directed evolution assessment library; this may arise because cysteines form disulfide bonds and can facilitate the protrusion of motifs on the surface of proteins^33^. While having a single cysteine in the insert sequence slightly elevates the median EV enrichment score *S* compared to no cysteines, more than one cysteine reduces fitness (Supplementary Fig. S7). In addition, we show that cysteines do not solely explain the improvement in manufacturing efficiency achieved through the ML model.

Finally, we investigated whether training with data from the ML assessment library would lead to a model performance improvement (Supplementary Fig. S2b and S3b). When trained on the ML assessment library, our LSTM model with two hidden layers of 160 and 20 nodes and one densely connected layer with one node (referred to as LSTM(160,20)_Dense(1)) showed the highest average performance across cross-validation splits. This model outperformed the equivalent model trained on Round_3, with a Pearson’s correlation coefficient of 0.69 compared to 0.64, suggesting that iterative rounds of ML-assisted selection might be desirable for related protein engineering tasks.

### Optimizing data set generation

As demonstrated above, dataset collection and curation (for example, the number of selection cycles performed) exhibited a stronger influence on model performance compared to model architecture, motivating the hypothesis that optimising dataset generation will yield larger gains in predictive accuracy than further architectural refinement (Fig.1d). To investigate which experimental factors most strongly constrain model performance, we conducted a series of controlled *in silico* perturbations, where we independently varied dataset volume, the size of amino acid sequence space explored, sequencing read depth and measurement noise.

Simulating a reduction in data volume showed that decreasing the total number of data points (i.e., the number of amino acid sequences screened) led to a strong decline in performance followed by a plateau, suggesting that data volume is the main parameter to tweak for gaining performance gains (Fig. 4a). We compared predictive performance across different amino acid sequence spaces (8-mer, 9-mer, 12-mer, and 13-mer), while controlling for diverging dataset volumes, and observed a decrease in ML model performance as the number of inserted amino acids increased (Fig. 4b). Datasets for 9-mer and 13-mer mutants were less diverse than for 8-mer and 12-mer mutants, explaining the dip in LR model performance. Interestingly, more complex model architectures (FNN, CNN, LSTM) were more robust to lower dataset diversity. Simulating a reduction in sequence depth revealed that choosing a more economical read coverage of 120K compared to 12 million reads (representing a 100-fold read count reduction) would result in a drop in model performance from 0.615 to 0.502 (Fig. 4c). Finally, increasing the level of Gaussian noise added to the log-transformed enrichment score *S* caused a steady reduction in predictive accuracy (Fig. 4d).

**Fig. 4:**
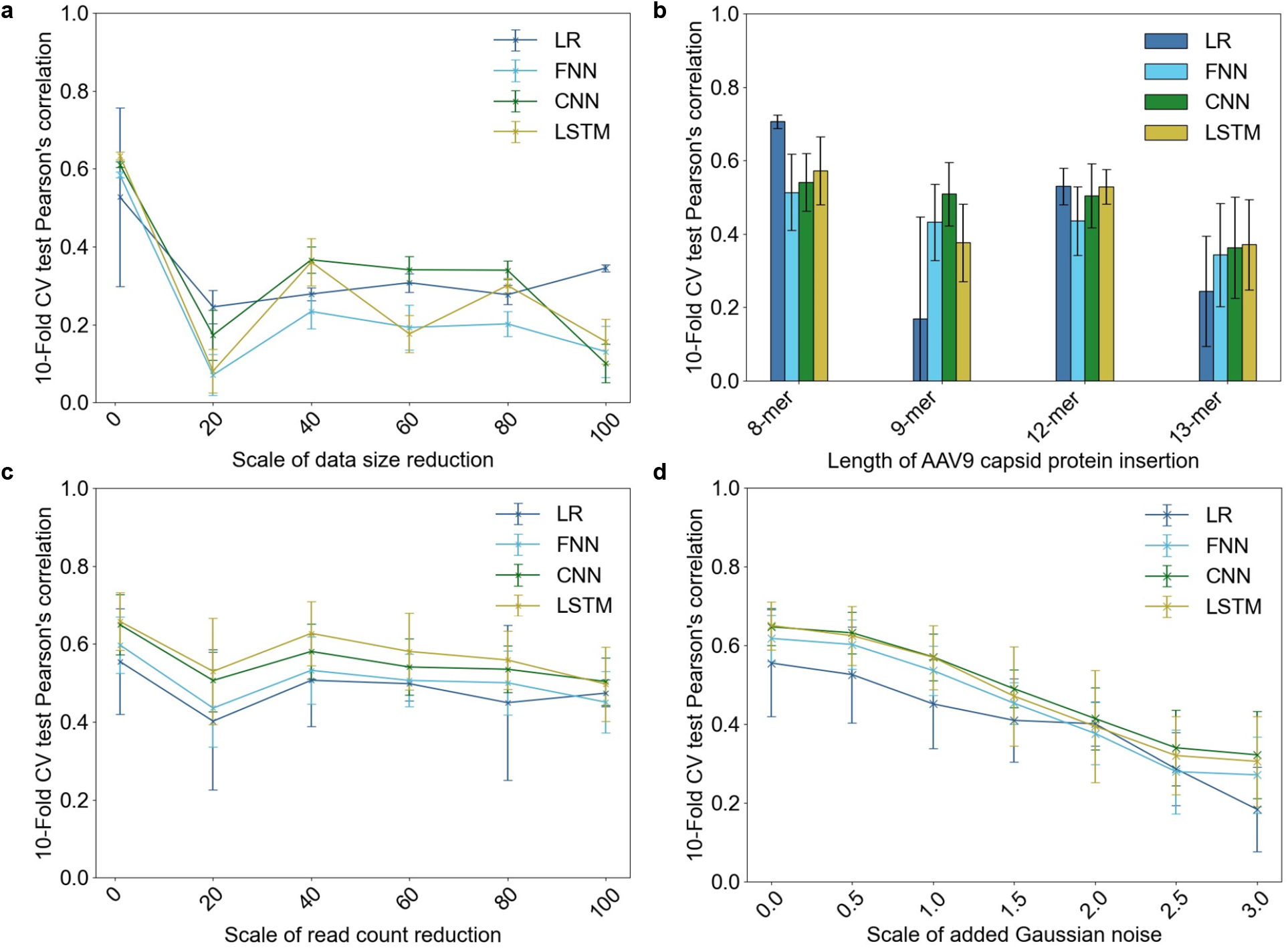
Analysing experimental parameters that influence ML model performance. Linear regression (LR), feed-forward neural network (FNN), convolutional neural network and long short-term memory (LSTM) models were trained to predict the log-transformed enrichment score *S* (an estimate for EV-AAV manufacturing efficiency) from the one-hot encoded amino acid sequence that was inserted into the AAV capsid protein. Model performance was measured in Pearson’s correlation on a held-out test set under 10-fold cross-validation (CV). (**a**) To investigate the influence of data volume on model performance, we divided the number of training data points (i.e., sequence-to-function relationships) by different factors, ranging from 1 (whole dataset) to 100 (99% of data points discarded). (**b**) To investigate the impact of the design space on model performance, training data points were filtered based on the number of amino acids in the insert sequence. (**c**) Read counts per mutant were divided by different factors before calculating the enrichment score *S*, demonstrating the influence of the read depth on ML model performance. (**d**) Model performance was plotted following the introduction of Gaussian noise parameterised by the coefficient of variation (standard deviation divided by mean fitness). For reference, the measured experimental coefficient of variation was approximately 0.5. Error bars represent ± standard error.

## Discussion

In this study, we benchmarked ML-assisted design against directed evolution on a novel AAV capsid engineering task. We show that integrating ML into directed evolution workflows can achieve a 15-fold improvement in mutant fitness, highlighting the potential of this approach for protein engineering. This experimentally validated gain is particularly remarkable in the light of ongoing concerns about model overfitting and the limited generalisability of ML efforts in the biosciences^34^.

AAV capsids have been engineered previously to increase AAV titres^14^, antibody evasion^7^, and tissue specificity^35^. Here, we instead optimised the encapsulation of AAVs into EVs, an approach shown to reduce immunogenicity and increase transduction efficiency in cultured cells^36^ and mice^37–39^. Despite the added technical complexity of separating EV-associated AAVs from conventional AAVs, our model outperformed previously published models trained to predict AAV capsid sequences associated with high AAV titres^14^. Model performance could potentially be further improved by using protein language models to create embeddings, as demonstrated by Wu et al., who combined protein language model embeddings with classical ML to achieve high Pearson’s correlations for AAV2 capsid fitness (*r*=0.818)^16^.

Our model tuning results indicate that training data quality (in terms of the number of selection rounds and sequencing reads per mutant) was the principal determinant of model performance, outweighing the choice of model architecture or hyperparameters. After a single round of selection, predictive performance was low (*r*=0.115, Supplementary Fig. S1), as measurement noise concealed true changes in mutant enrichment. In contrast, after *three* rounds of selection, enrichment of high-performing mutants exceeded the experimental noise floor (*r*>0.6, Supplementary Fig. S3a). Meanwhile, model architecture did not have a significant impact on model performance. This observation is corroborated by Silverstein et al., who found comparable performance across neural networks, linear regression and random forest models on an ML-assisted CRISPR-Cas9 PAM engineering task^40^. It also aligns with findings from Yang et al., who showed that CNNs perform similarly to transformers on protein prediction tasks^30^ and from Bryant et al., who showed that dataset composition matters more than architecture for predicting AAV capsid viability ^15^.

Our results suggest that ML model performance – and, hence, AAV candidate selection – depends on the unique fingerprint of each protein engineering dataset. Optimising experimental design parameters is critical for generating effective training data but necessitates balancing throughput and data quality. In our case, this balance involved trade-offs between library size, measurement throughput, sequencing depth, measurement noise and the number of selection rounds (each of which enriches mutants above background fluctuations). As such, it is desirable to have a framework for selecting these experimental parameters for guiding data set generation, which remains the primary bottleneck in terms of resource and time requirements for work such as ours (see also that in Vorobieva *et al*^*25*^).

Initial efforts have begun to examine the “ruggedness” of the fitness landscape as a parameter involved in the success of protein engineering efforts. For example, Li et al. analysed sixteen fitness landscapes and showed that ML-assisted directed evolution performance varied substantially across landscapes^5^. Similarly, the data volume and quality required for generalisable ML-assisted antibody binding have been investigated^41^. However, these studies are neither exhaustive nor sufficient for guiding experimental design across diverse engineering tasks. Future dataset generation strategies should therefore be optimised jointly with ML experts, so that experimental design, model development and candidate selection are treated as an integrated workflow.

Future research could also investigate *what* ML models learn about protein function. Explainability techniques have been studied to interpret attention in protein language models, but not specifically in the ML-assisted directed evolution context. An additional hurdle is finding explanations that are specific to the protein engineering task at hand, given the broad applicability of protein language models^42^. While there is a plethora of interpretability methods within ML, applying them to uncover biological explanations remains challenging^43,44^. Future work could also apply transfer learning methods to train models on previously published AAV data and fine-tune on EV-encapsulation data. In addition, models might be trained to optimise multiple objectives simultaneously (e.g., one on AAV viability data and another on EV recruitment), with predictions aggregated to identify candidate capsids with favourable combined properties.

By closing much of the manufacturing gap between EV-AAV and standard AAV vectors, this work removes a major bottleneck that has constrained the development of EV-AAV as a gene therapy platform, and opens up a clear path towards its clinical translation. Building on this manufacturing advance, a natural next step is to characterise the tissue tropism and biodistribution of the evolved capsids, which would clarify the therapeutic doses achievable with these higher-yielding variants and, given the reduced immunogenicity and improved transduction efficiency already associated with EV encapsulation^21^, may support dose de-escalation relative to standard AAV, an important consideration given the safety concerns associated with high systemic AAV doses^6^. Validating capsid performance in higher-order animal models and human tissue would further strengthen the case for clinical translation, building on precedents for cross-species validation of engineered AAV capsids^8^.

Another approach to target the multi-faceted design challenge that AAV engineering presents is to use our model as a tool for selecting manufacturable candidates from screens aimed at other clinically relevant properties (e.g., bioavailability) or combine them with models trained to predict such properties. Moreover, tissue tropism could be conferred *post-hoc* by fusing tissue-specific targeting peptides to exosomal membrane proteins (as previously shown by Alvarez-Erviti et al.^45^).

To summarise, our work has identified AAV capsid mutants with up to 41-fold improved overall EV-AAV manufacturing efficiency, and mutants with up to 99.2% encapsulation efficiency, representing a substantial advance over previous approaches and a step towards closing the manufacturing yield gap between EV-AAV and standard AAV gene therapy vectors. Our work also provides strong quantitative evidence (highlighted by a >15-fold performance improvement) that even simple ML methods with moderate predictive performance can interpolate between observed data points and design libraries that outperform those based solely on selecting experimentally observed winners. Given the massive protein design space (e.g., 25.6 billion possible mutants for an 8-mer insertion), the probability of identifying an equivalently high-performing mutant through screening alone is low. As such, application of our method to increase the likelihood of selecting a top candidate via learning from initial rounds of screening – which here required no experimental work beyond that already performed for directed evolution – promises to be an effective enabler of future bioengineering strategies.

## Methods

### AAV capsid library synthesis

We created libraries by inserting variable eight, nine, twelve or thirteen amino acids (8-, 9-, 12- or 13-mers) in position 453-454 of the AAV9 capsid. Therefore, single-stranded (NNK)_8-13_ oligos with BsmBI-v2 compatible overhangs were synthesised (IDT) and converted to double-stranded DNA using Klenow fragment–mediated extension. Next, the DNA fragment was digested with BsmBI-v2, purified by PCR cleanup and quantified by NanoDrop spectrophotometry. For plasmid assembly, replication-incompetent AAV9 plasmid backbone was first digested with MLuI to prevent re-ligation, followed by BsmBI-v2, according to manufacturer protocols (New England Biolabs). Digested plasmids were treated with Antarctic phosphatase at 37 °C for 30 min and heat-inactivated at 80 °C for 2 min. Subsequently, backbone fragments were purified either by gel extraction or PCR cleanup, depending on band quality, and quantified by NanoDrop spectrophotometry. Insert and backbone fragments were assembled using Gibson Assembly Master Mix (New England Biolabs), yielding plasmid libraries. Each 20 µL reaction contained 10 µL Master Mix, 50 ng plasmid backbone, and 3.4 ng insert DNA, with nuclease-free water added to volume. Assemblies were incubated at 50 °C for 1 h.

NNK oligos were used to reduce the fraction of amino acid inserts containing stop codons, which lowers the proportion of 8-mers carrying at least one stop from 32% under fully degenerate NNN to 22%. To assess whether this came at the cost of a skewed amino acid representation, we calculated the expected per-position amino acid frequencies of both schemes from the standard genetic code. Aside from tryptophan and methionine (raised from 1.6% to 3.1% per position) and isoleucine (lowered from 4.7% to 3.1%), the two schemes yield identical frequencies. Judged against a uniform amino acid distribution, these shifts leave NNK marginally closer to uniform than NNN (summed absolute deviation 0.426 versus 0.446, stop codons excluded), and neither scheme approximates the amino acid composition of the wild-type AAV9 capsid protein more closely than the other.

### Pre-encapsulation library isolation

Following the generation of plasmid libraries that encode AAV capsid mutants, libraries had to be amplified to reach the required yields for transfection into HEK 293 T cells. Therefore, assembled plasmid libraries were electroporated into electrocompetent *Escherichia coli* (New England Biolabs, C3020K) at ≤10% of the cell volume to minimize toxicity. Cells were recovered in pre-warmed SOC Outgrowth Medium (New England Biolabs) within 30 s. After recovery, 40 µL of the electroporated cells were plated onto 15 cm LB-agar plates containing ampicillin and spread evenly. Electroporation efficiency and library diversity were estimated by plating a 1:1000 dilution and counting colony-forming units. Colonies were harvested by gently scraping plates with cold LB (2–5 mL per plate) and pooling into pre-chilled 50 mL tubes. Cells were pelleted by centrifugation at 4 °C (4,000 g, 15–20 min), and supernatants were discarded. Finally, plasmid DNA was purified using commercial Maxi Kits (Qiagen or Macherey-Nagel), yielding pre-encapsulation libraries.

### EV-AAV and AAV production

To produce EV-AAVs and AAVs from plasmids, libraries were transfected into HEK 293 T cells. Cell culture procedures involved maintaining HEK 293 T cells in Dulbecco’s Modified Eagle Medium supplemented with GlutaMAX (Gibco), 10% fetal bovine serum and 1% penicillin/streptomycin. Cells were kept at 80-90% confluency by being harvested from T175 flasks using trypsinization, centrifuged at 400 g for 5 min, and resuspended in fresh growth medium. For transfection, cells were seeded onto 15 cm tissue culture dishes at densities adjusted to reach 70–80% confluency at the time of transfection. Plasmid transfections were performed using PEImax (ThermoFisher). For each 15 cm dish, pHelper and pRepHelper at equimolar ratios, and 100 ng of the pre-encapsulation library were combined. Plasmid DNA was diluted in Opti-MEM (ThermoFisher) to a final volume of 1 mL per plate. In parallel, PEImax (10 mg/mL stock) was diluted in Opti-MEM at a 3:1 mass ratio relative to DNA. PEImax and DNA solutions were combined dropwise and incubated for at least 20 min at room temperature to allow complex formation. Complexes (2 mL per plate) were added dropwise to cells and distributed evenly before returning the dishes to the incubator. Four hours after transfection, media was replaced with 20 mL Opti-MEM supplemented with 1% penicillin/streptomycin per dish. During the following incubation period, AAV and EV-AAV particles are formed and released from the cells into the supernatant.

### EV-AAV and AAV purification

Supernatant containing secreted EV-AAVs and AAVs was harvested 48-72 h post-transfection. Media from multiple dishes were pooled and purified by sequential centrifugation at 400 g for 5 min followed by 4,000 g for 25 min to remove cells and debris. The supernatant was concentrated using tangential flow filtration followed by centrifugal ultrafiltration. Samples were concentrated with intermittent dilution in PBS to prevent viscosity-related precipitation. Final concentration was achieved using centrifugal filters with 100 kDa molecular weight cutoff at 4 °C.

To measure intracellular AAV capsid sequences, AAV genomes that were not released into the supernatant were isolated. Therefore, HEK 293 T cell pellets were lysed in 50 mM tris and 150 mM NaCl (pH 8.5) buffer and subjected to four freeze/thaw cycles in dry ice/ethanol. Thawed lysates were supplemented with Benzonase to a final concentration of 50 U/ml and incubated for 30–45 min at 37 °C. Lysates were purified by centrifugation at 3700 g for 20 min at room temperature and the supernatant was passed through a 0.45 μm syringe filter.

Next, EV-AAVs and AAVs were isolated from supernatant and lysate samples via iodixanol density centrifugation. Iodixanol gradients were prepared by sequential layering of defined iodixanol solutions, starting from the highest density fraction. Concentrated samples (500 µL) were layered onto the gradients and fractionated using ultracentrifugation. Fractions corresponding to EV-AAVs (typically fractions 4–6) or AAVs (fractions 9–10) were collected, diluted in PBS and concentrated using centrifugal filters with 100 kDa molecular weight cutoff. Multiple wash steps were performed until iodixanol was eliminated. Final purified EV-AAV and AAV preparations were concentrated to 250–500 µL, aliquoted (50–100 µL and stored at −80 °C.

### Post-encapsulation library isolation

To isolate encapsulated AAV genomes, EV-AAV and AAV preparations were first treated with DNase (Promega) to degrade surface-bound DNA. Reactions were assembled in PCR tubes to ensure accurate temperature control: 10× DNase buffer, 1 µL DNase, 5µL EV-AAV or AAV sample and nuclease-free water to a final volume of 17 µL. Samples were incubated at 37 °C for 30 min, followed by DNase inactivation with 2 µL stop solution and incubation at 65 °C for 10 min. Next, reactions were cooled on ice. To open up capsids and lipid bilayers, preparations were treated with PCR-grade proteinase K (Roche), revealing encapsulated AAV genomes. A digestion mix containing 2× proteinase K buffer (0.1 M EDTA, 0.5% SDS), proteinase K, and nuclease-free water was prepared, and 180 µL was added to each DNase-treated sample. Samples were incubated at 50 °C for 1 h and then cooled on ice. For next-generation sequencing, the variable insert region of the isolated AAV genomes was amplified using high-fidelity polymerase (KAPA HiFi, Roche) and flanking primers. Amplicons were purified using Monarch kits and submitted for sequencing (Azenta or BGI).

### Next-generation sequencing data processing

*Python* (version 3.7.12) and *bash* (version 5.1.16) scripts were used to process raw FASTQ files. FASTQ files corresponding to the same sample were identified based on the sample ID and concatenated into a single FASTQ file. Next, paired-end reads were merged. For each read pair, reverse reads were reverse-complemented and aligned against the forward read by sliding overlap alignment, with reads processed in parallel using multiprocessing. The optimal overlap was selected by maximizing the number of matching bases, allowing ambiguous nucleotides to match any base. Successfully aligned pairs were merged into a single sequence, while read pairs with failed alignments (and, hence, of low read quality) were discarded. Subsequently, merged reads were filtered to retain only sequences containing both the left and right constant regions that flank the variable insert, following the format: 5’-TATCTCTCAAAGACTATTAAC-(NNK)_8-13_-(ACGCTAAAATTCAGTGTGGCC)-3’.

This filtering step was performed using linked adapter identification with *cutadapt* (version 2.6)^46^, allowing partial mismatches (based on the default error rate of 10%) while requiring a minimum overlap length of 21 nucleotides for each flanking region. The flanking regions were then trimmed to extract the 24 to 39 nucleotide insertions (corresponding to between 8 and 13 variable amino acids). Filtered FASTQ files were parsed using *biopython* (version 1.81) and converted into dataframes using *pandas* (version 1.3.5). The resulting dataframes contained columns for read ID, nucleotide sequence and Phred quality score. Nucleotide sequence lengths were computed for all reads and only sequences within the expected length range were retained. Lastly, AAV capsid mutants were grouped by nucleotide sequence identity, and counts per unique mutant were computed. Sequencing recovered 24,169 distinct mutants in the pre-encapsulation library and 26,901 in the post-encapsulation library, of which 18,597 were present in both and therefore retained for the calculation of S.

### Data normalization

Mutant counts were normalized by the sequencing read depth per sample with:

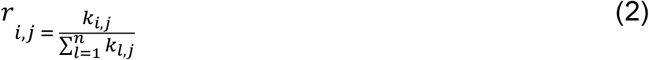

Where *r* is the normalized mutant count, *k* is the raw count for *i* = 1, …, *n* sequences and *j* = 1, …, *m* samples.

Next, the enrichment score *S* (as estimate for the EV-AAV manufacturing efficiency) was calculated by comparing normalized mutant count frequencies between pre- and post-encapsulation libraries:

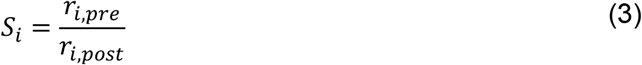

Where *r* is the normalized mutant count in the pre-encapsulation and post-encapsulation library. To avoid dividing by 0 when calculating *S*, only mutants with read counts in both the pre-encapsulated and post-encapsulated libraries were retained (except for training Round_3_Pseudo_count, where missing sequences were assigned a read count of 1 following Equation 4).

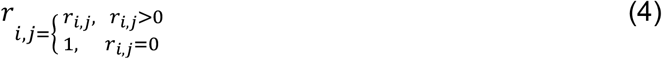

Finally, the enrichment scores *S* were aggregated across codon replicates *c* (i.e., nucleotide sequences that encode for the same amino acid sequence).

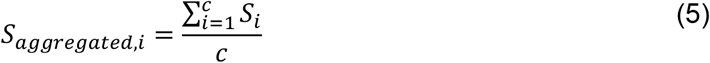

### ML model training and evaluation

Amino acid sequences and corresponding enrichment scores *S* were pre-processed using *Python* scripts (version 3.12.5). Specifically, amino acid sequences were one-hot encoded into a 20 × *l* matrix, where *l* represents the number of amino acids (e.g., an 8-mer yields a 20 × 8 matrix). Enrichment scores *S* were log-transformed to place enrichment and depletion on a symmetric scale: as a ratio, *S* is bounded below by zero and unbounded above, so untransformed values compress depletion into the interval [0, 1] while enrichment spans [1, ∞). For training, the dataset was randomly shuffled and split into training, validation and test sets (containing 80%, 10% and 10% of the data, respectively). We trained models using datasets generated in the first and third round of selection, which will be referred to as Round_1 and Round_3.

We also applied three different pre-processing techniques. Two of these pre-processing techniques take into account the statistical power of *S* (which is stronger when the mutant counts *r*_*i,j*_ in the pre-encapsulation and post-encapsulation libraries are higher). One of these strategies is discarding mutants with mutant counts that are lower than a threshold. We evaluated which threshold would discard 90% of mutant sequences with stop codons, which should not form viable capsid proteins, but can occur in the post-encapsulation library due to cross-packaging. The other strategy is to compute the variance of *S* using Equation 6, following an approach published previously^47^, and weight each sequence by training the model with a weighted loss ℒ (Equation 7). Here, *M* represents the number of data points, *y*_*i*_ the output variable, *x*_*i*_ the input variable and *f* the predictive model.

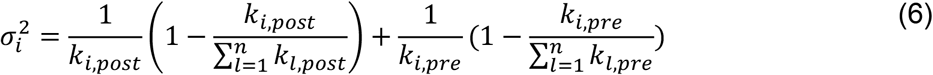

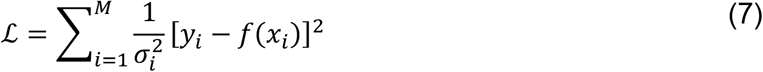

The third pre-processing strategy involves applying a pseudo count of 1 to mutants missing in either the pre-encapsulation or post-encapsulation library (see Data Normalization above).

To assess whether predictive performance depended on sequence similarity between splits, we additionally generated similarity-aware splits. For each identity cutoff – 87.5%, 75%, 62.5% and 50%, corresponding to one to four mismatches across the 8-mer – test sequences were required to fall below that identity to every training sequence, with sequences violating the constraint reassigned to the training set. Because identity between 8-mers is quantised in 12.5% increments, these represent the full set of available thresholds above 50%. Predictive performance was evaluated as Pearson’s *r* on held-out test data under 10-fold cross-validation.

For function *f*, we compared linear regression (LR) with feed-forward neural networks (FNNs), convolutional neural networks (CNNs) and long short-term memory (LSTM) models. LR is used as a baseline due to its excellent interpretability (each amino acid at each position in the insert gets assigned a weight and, hence, high-fitness amino acids can be easily extracted). Neural networks (such as FNNs, CNNs, and LSTMs) can describe non-linear, more complex relationships between *x*_*i*_ and *y*_*i*_ but are prone to overfitting (performing well on the test set but not on a separate assessment library). Linear models were implemented in *scikit-learn* (version 1.5.2) and neural networks using *Keras* in *TensorFlow* (version 3.6.0). In total, we compared fifteen models: five LSTMs with one, two and three hidden layers of 20 and 160 nodes, with one or two dense layers of eight or one nodes which were trained on Round_3 and the ML assessment library; two CNN and two FNN models with two hidden layers of 160 nodes and one dense layer which were trained on Round_3 and the ML assessment library; and six linear models trained on Round_1, Round_3 and the ML assessment library with the different pre-processing strategies outlined above. To compare model performance, we calculated Pearson’s correlation between predicted and observed values on a held-out test set under five-fold cross-validation.

Unless stated otherwise, performance is reported on the complete held-out test set. For the sensitivity analysis, data were randomly shuffled and split as above, before any ranking by enrichment score. The resulting test set was then sorted by measured enrichment score and truncated sequentially to retain the top 90%, 80%, …, 20% of mutants, discarding the lowest-scoring remainder at each step. Training sets were left unchanged by this procedure.

### ML-assisted library design

To design novel amino acid sequences using Round_3_LR_Thresholding, we loaded models from five independent train-validate-test folds and averaged model weights *w*_*p,a*_ across folds (see Equation 1). Model weights cannot function as probabilities since they are not restricted to the interval [0,1] and roughly half are negative. We decided to use reciprocal-rank transform for mapping ℝ → ℝ_+_ monotonically because residue *order* was reproducible across the five folds while the absolute weights were not, and because it prevents any single residue from dominating a position and collapsing the library onto one consensus sequence. Therefore, we assigned a zero-indexed rank *rk*_*p,a*_ to each amino acid *a* per insert position *p* in descending order of their average *w*_*p,a*_ and converted *rk*_*p,a*_ to reciprocal-rank weights *R*_*i,a*_ (see Equation 8). Reciprocal-rank weights *R*_*i,a*_ were normalised to give a categorical probability distribution over the amino acid alphabet *P*(*a*|*p*) by dividing *R*_*p,a*_ of each amino acid by the sum of the reciprocal-rank weights of all 20 amino acids at that position (see Equation 9). A total of 472 novel amino acid sequences were designed by drawing each of the eight positions independently from its corresponding distribution *P*(*a*|*p*)).

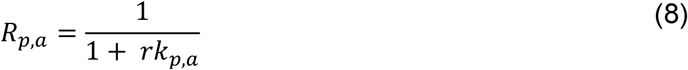

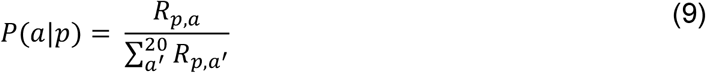

### Parameter ablation studies

We investigated the influence of different experimental parameters (dataset volume, read coverage, amino acid design space and measurement noise) on ML model performance. Therefore, we used simulated datasets based on Round_3 to train LR, FNN(160, 160)_Dense(1), CNN(160)_Dense(1) and LSTM(160, 20)_Dense(1) models, applying a read count threshold of 32 (see Data normalization). First, we ablated the number of datapoints in the training data by dividing the total number (18K) by varying divisors (from 20 to 100), keeping the size of the test set constant. Second, we trained ML models on datasets with varying amino acid insert lengths (i.e., changing the amino acid design space investigated). To obtain enough data points for model training, we included rationally designed sequences from previous work^25^. Third, we simulated datasets with reduced read coverage by dividing the mutant count in the pre-encapsulation and post-encapsulation libraries by varying read count divisors (from 20 to 100). Non-integer values generated during read count division were converted into whole numbers using probabilistic rounding. Specifically, each value was rounded either up or down based on the remainder when the read count could not be divided exactly by the read count divisor. The probability of rounding upwards was equal to the decimal fraction of the original non-integer value, preserving the underlying distribution while minimizing systematic rounding errors. Finally, we simulated datasets with increased measurement noise by randomly drawing enrichment scores *S* from a Gaussian distribution *p*(*S*) with varying standard deviations *σ* (see Equation 8). Hereby, standard deviations *σ* were selected as a fraction or multiple of the mean *µ* (ranging from 0.5 to 3.0).

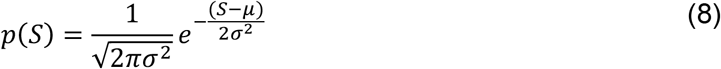

## Authorship information

This study was conceptualised and supervised by M.L. and H.S., with input from D.G. and S.J.S. Its methods were determined by M.L. and I.V. The investigation was conducted by M.L., R.M., M.J., and S.C. Formal analysis and visualisation were performed by M.L. Funding for this study was acquired by M.J.A.W. The original draft was written by M.L. and reviewed and edited by H.S. and M.L.

D.G. is a stakeholder in Evox Therapeutic Ltd (UK), M.J.A.W. is a consultant and stakeholder in Evox Therapeutic Ltd (UK). All other co-authors reported no conflict of interest.

## Acknowledgements

We thank the members of the Steel, Sanders and Wood laboratory, specifically Hans Friedrichsen, Sebastian Towers, Emilie Wigdor, Olivia Gallupova and Narjes Rohani for continuous, extensive discussions of the project. Additionally, we thank Mariana Conceicao of Evox Therapeutics for discussions. M.L. received funding from the Interdisciplinary Bioscience Doctoral Training Partnership at the University of Oxford and UK Research and Innovation (BB/T008784/1). I.V. was supported by a PhD studentship from Evox Therapeutics. D.G. is supported by a Hjärnfonden Postdoctoral Fellowship, CureHeart, the British Heart Foundation’s Big Beat Challenge Award (BBC/F/21/220106), BBSRC New Investigator Award (UKRI2992) and MRC Core in Therapeutic genomics (MR/Z504725/1). This work was additionally supported by the Cystic Fibrosis Foundation (CFF; HNR02540), the U.S. Department of Defence (DoD; A3R00010; W81XWH-22-1-1067), and Muscular Dystrophy UK (MDUK; HNR01740; 19GRO-PG36-0339).

## Data availability

NGS data generated in the study that are needed to reproduce, verify, and extend the findings are available at the NCBI SRA under accession: PRJNA1473000. There are no restrictions on data availability.

## Code availability

Code produced in the study that is needed to reproduce, verify, and extend the findings of the study is available and will be maintained in the GitHub repository: https://github.com/marinaluchner/ml-assisted-aav-engineering.

