## Supplemental Data 1 for "Machine learning-assisted directed evolution yields dramatic improvement on novel AAV engineering task"

**Contents:**

Supplementary Figures S1 to S7 and Table S1.

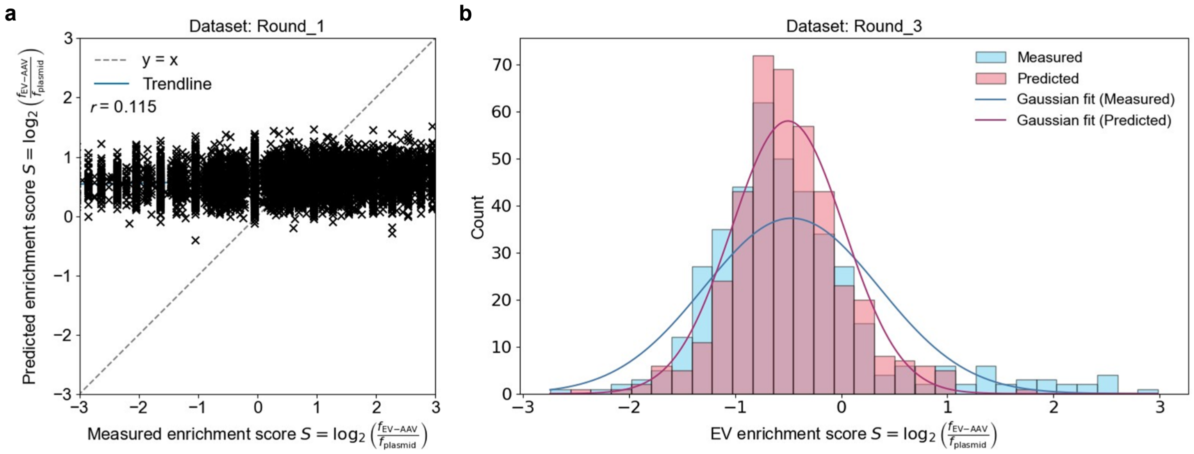


**Supplementary Fig. S1: Tuning ML model performance to learn EV recruitment of AAV9 from amino acid sequence.** (**a**) Correlation between ML model predictions and experimentally measured ("true") log enrichment scores $S$, from data derived during the first round of selection, excluding $S$ by thresholding the read count (Round_1_LR_Threshold). (**b**) Overlap of predicted and measured log enrichment score $S$ distributions from data generated in the third round of selection (Round_3).

**Supplementary Table S1: ML models trained to predict the log EV enrichment score from one-hot encoded AAV9 capsid insert sequences.** Each row represents one model class, defined as a unique combination of architecture, hyperparameters and training dataset; models sharing an architecture but differing in training data or hyperparameters are counted separately. Bracketed values give the number of nodes in each successive hidden and dense layer. "Thresholded" denotes application of a read-count threshold, discarding 90% of mutant sequences containing stop codons. Pearson's *r* is the average on a held-out test set under five-fold cross-validation.

| Model | Training data | Architecture | Hyperparameter | Pearson's r |
|---|---|---|---|---|
| **Linear regression (LR)** | | | | |
| Round_1_LR | Round_1 | LR | Weight $w$ per (insert position $p$, amino acid $a$) pair plus bias $b$; one-hot input; no pre-processing | 0.124 |
| Round_3_LR | Round_3 | LR | As above; no pre-processing | 0.354 |
| Round_3_LR_Weight | Round_3 | LR | As above; loss weighted by $\frac{1}{\sigma^2}$ of $S$ (Eq. 6–7) | 0.363 |
| Round_3_LR_Threshold | Round_3 | LR | As above; read-count threshold to discard 90% of mutant sequences with stop codons | 0.611 |

| Model | Training data | Architecture | Hyperparameter | Pearson's r |
|---|---|---|---|---|
| Round_3_LR_Pseudo_ count | Round_3 | LR | As above; missing read counts replaced by pseudo count of 1 (Eq. 4) | 0.390 |
| ML_Assessment_Library _LR_Threshold | ML_Assessment_ Library (thresholded) | LR | As above; read-count threshold to discard 90% of mutant sequences with stop codons | 0.568 |
| **Long short-term memory (LSTM)** | | | | |
| LSTM(20)_Dense(1) | Round_3 (thresholded) | LSTM | 1 hidden layer, 20 nodes; 1 dense layer, 1 node | 0.625 |
| LSTM(160,20)_Dense(1) | Round_3 (thresholded) | LSTM | 2 hidden layers, 160 and 20 nodes; 1 dense layer, 1 node | 0.650 |
| LSTM(160,160,160)_ Dense(1) | Round_3 (thresholded) | LSTM | 3 hidden layers, 160 nodes each; 1 dense layer, 1 node | 0.646 |
| LSTM(160,20)_ Dense(8,1) | Round_3 (thresholded) | LSTM | 2 hidden layers, 160 and 20 nodes; 2 dense layers, 8 and 1 nodes | 0.648 |
| LSTM(160,20)_Dense(1) | ML_Assessment_ Library (thresholded) | LSTM | 2 hidden layers, 160 and 20 nodes; 1 dense layer, 1 node | 0.682 |
| **Feed-forward neural network (FNN)** | | | | |
| Dense(160,160,1) | Round_3 (thresholded) | FNN | 2 hidden layers, 160 nodes each; output layer, 1 node | 0.616 |

| Model | Training data | Architecture | Hyperparameter | Pearson's r |
|---|---|---|---|---|
| Dense(160,160,1) | ML_Assessment_ Library (thresholded) | FNN | As above | 0.666 |
| **Convolutional neural network (CNN)** | | | | |
| Conv2D(160)_Dense(1) | Round_3 (thresholded) | CNN | 1 convolutional layer, 160 filters; 1 dense layer, 1 node. | 0.655 |
| Conv2D(160)_Dense(1) | ML_Assessment_ Library (thresholded) | CNN | As above | 0.673 |

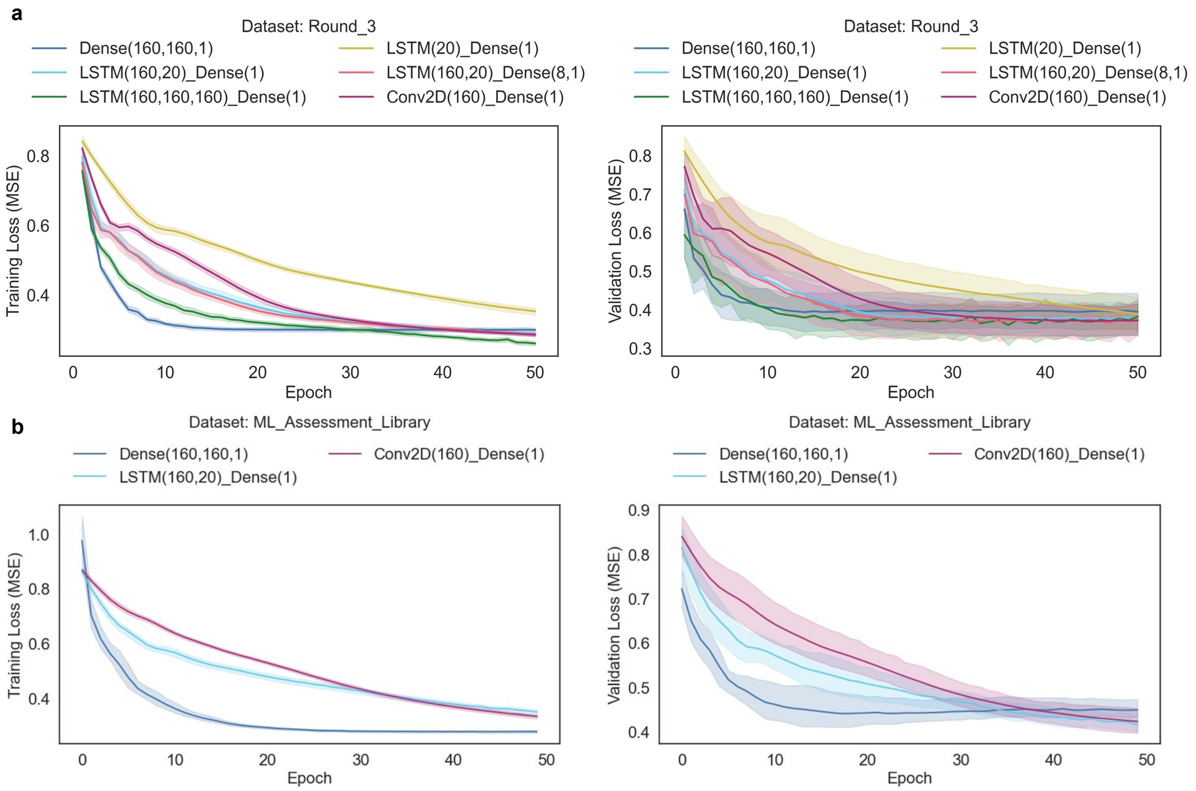


**Supplementary Fig. S2: Training and validation loss curves for the neural network architectures.** Mean squared error on the training set (left) and validation set (right) across 50 epochs for models trained on (**a**) the thresholded Round_3 dataset and (**b**) the thresholded ML assessment library. Lines give the mean across five cross-validation folds; shaded bands, ± confidence intervals. No loss curves are shown for the linear regression models, which are solved in closed form and therefore have no iterative training trajectory.

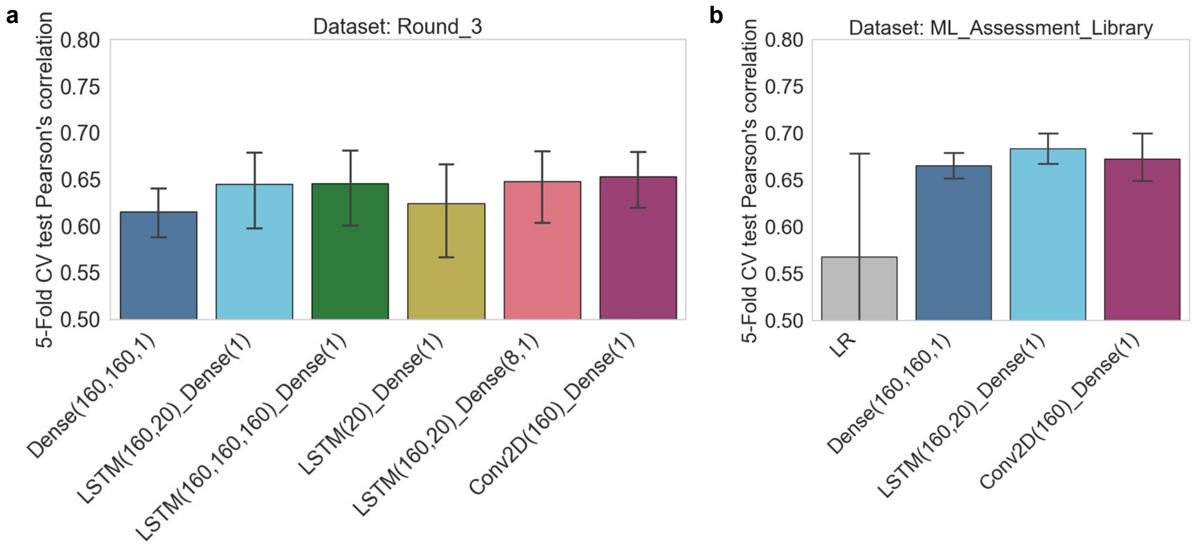


**Supplementary Fig. S3: Investigating the influence of model architecture and dataset design on model performance.** One-hot encoded amino acid sequences that were inserted into AAV capsid protein were used as input variables to predict the log enrichment score $S$ of these AAV capsid mutants (providing an estimate for their EV encapsulation efficiency). (**a**) Different models (LSTM, FNN, CNN) were tested using varying numbers of layers and nodes per layer, using data generated in the third round of selection (Round_3). (**b**) Re-training of these models and LR using data from the ML assessment screen presented in Fig. 3. Average Pearson's correlation under five-fold cross-validation (CV) $\pm$ 95% confidence intervals were plotted.

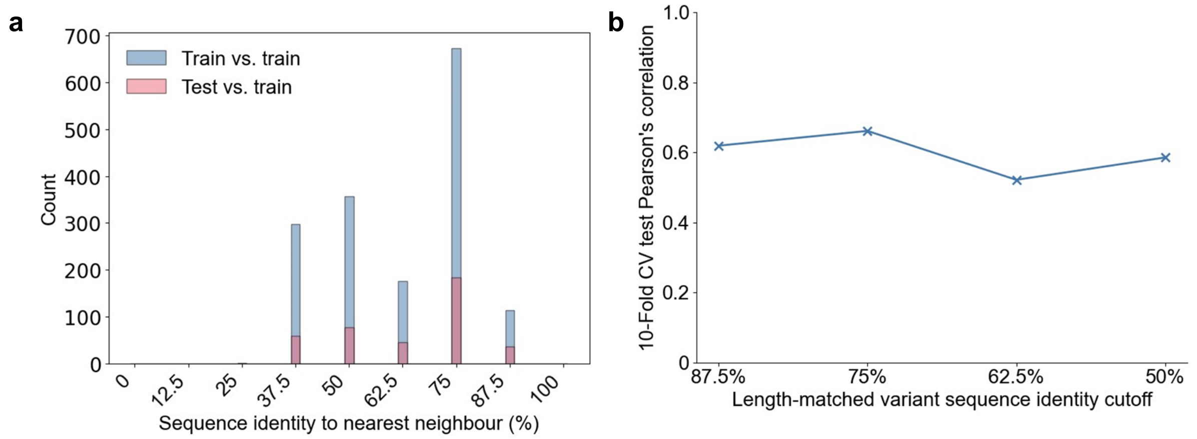


**Supplementary Fig. S4: Model performance is robust to similarity-aware data splitting.** (**a**) Distribution of nearest-neighbour sequence identity for the 8-mer inserts, comparing each training sequence with its closest other training sequence (blue) and each test sequence with its closest training sequence (red). No sequence exceeds 87.5% identity (one mismatch) to its nearest neighbour in the training set, and only 32 of 404 test sequences fall in this highest bin; 100% identity is impossible by construction, as every 8-mer occurs exactly once in the dataset. (**b**) Pearson's correlation coefficient on held-out test data under 10-fold cross-validation, with training and test sets split so that no test sequence exceeds the indicated length-matched sequence identity cutoff to any training sequence (87.5%, 75%, 62.5% and 50%, corresponding to 1-4 mismatches across the 8-mer insert).

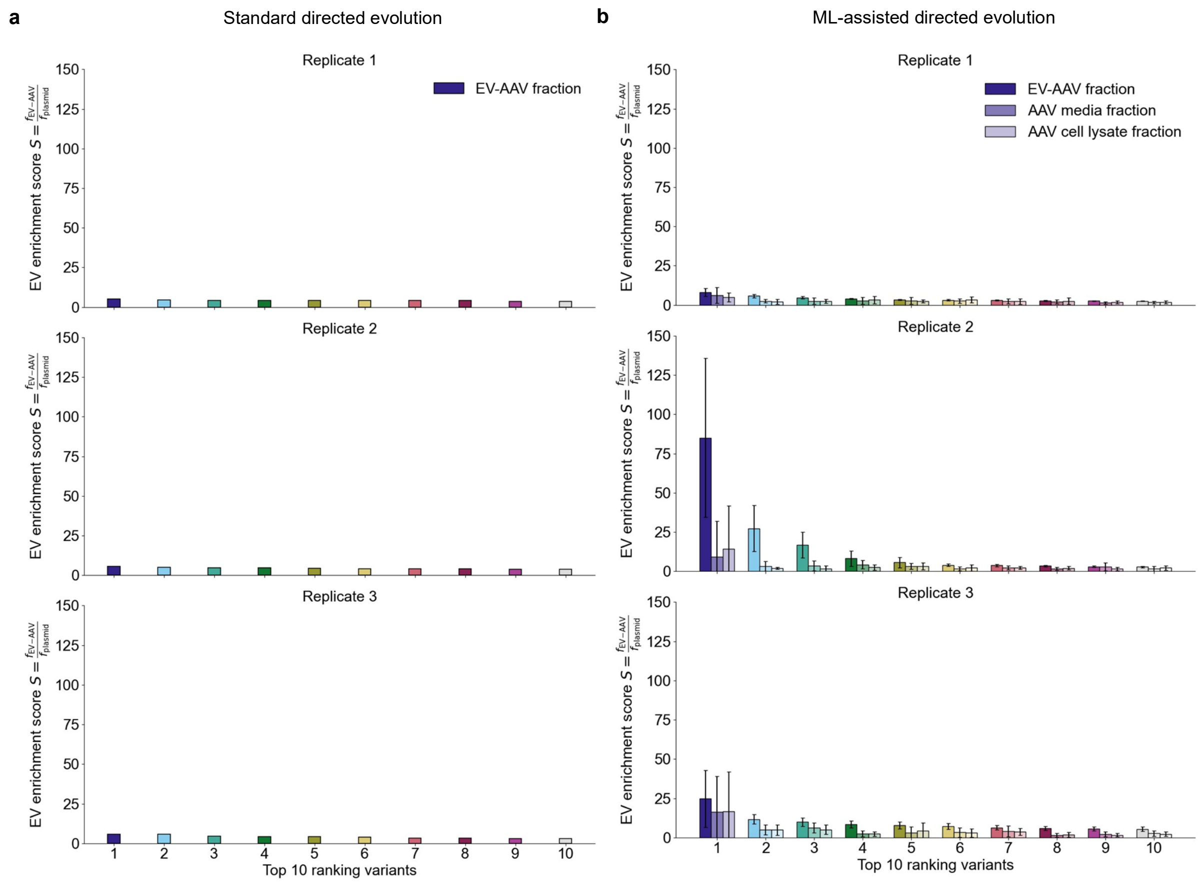


**Supplementary Fig. S5: Accounting for assessment library size when comparing standard against ML-assisted directed evolution.** Protein mutants were selected using either standard directed evolution or ML-assisted directed evolution. The EV encapsulation efficiency of the selected mutants was assessed by calculating the log enrichment score $S$, comparing the AAV genome counts in the post-encapsulation library to the pre-encapsulation library. (**a**) For the directed evolution assessment library, which comprised 123 mutants, EV enrichment scores $S$ were ranked and plotted for the top 10 ranking mutants. (**b**) As the ML assessment library comprised of 472 mutants, we randomly subsampled the library and ranked mutants based on their EV enrichment score $S$. The subsampling and subsequent ranking steps were performed ten times, and the average EV enrichment scores $S \pm$ standard error plotted.

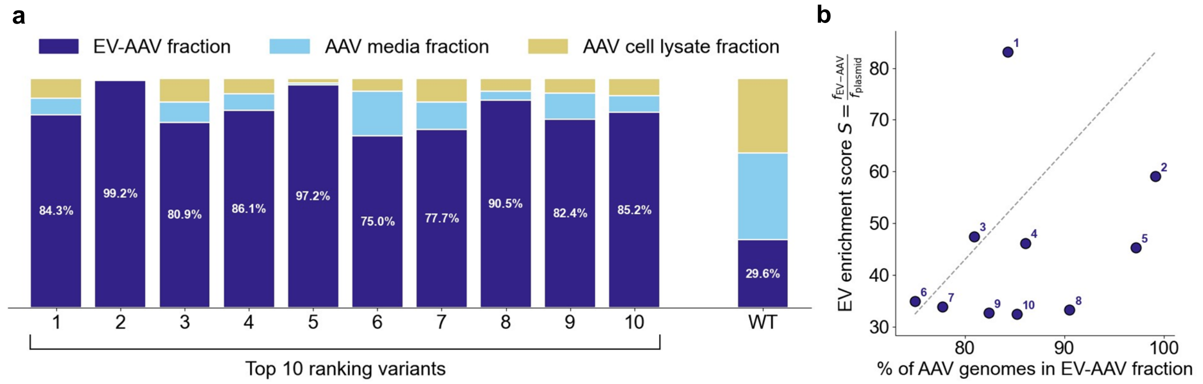


**Supplementary Fig. S6: EV encapsulation efficiency of the top-ranked ML-designed variants.** (**a**) Distribution of AAV genomes across the three purified fractions – EV-AAV (dark blue), AAV media (light blue) and AAV cell lysate (yellow) – for the top 10 ranking variants of the ML assessment screen and for wild-type (WT, GSGQNQQ). Values within the bars give the percentage of total AAV genomes recovered in the EV-AAV fraction. (**b**) EV enrichment score $S$ plotted against the percentage of AAV genomes in the EV-AAV fraction for the same 10 variants, illustrating that the two properties vary independently and can be used jointly to select variants with an optimal trade-off. The dashed grey line is a reference diagonal drawn between the minimum and maximum observed values of both metrics; variants lying close to this line show a balanced trade-off between the two properties, and those furthest towards the top right combine high EV enrichment with high encapsulation efficiency.

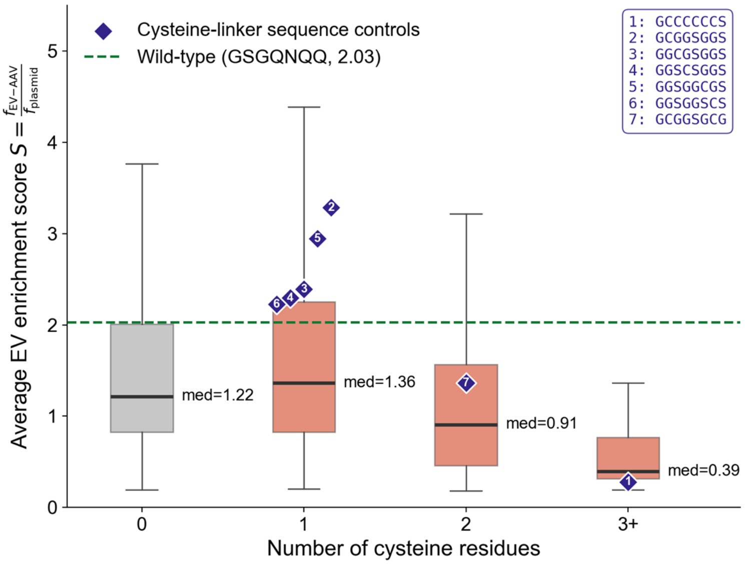


**Supplementary Fig. S7: Cysteine content alone does not explain the elevated EV enrichment of the top-ranked variants.** Average EV enrichment score $S$ for ML assessment library variants grouped by the number of cysteine residues in the 8-mer insert (0, 1, 2 or ≥3). Boxes show the interquartile range with the median indicated; whiskers extend to the 5th and 95th percentiles; medians are annotated. Blue diamonds mark the seven cysteine-linker sequence controls (sequences listed in the inset). Dashed green line, wild-type performance (GSGQNQQ, $S$ = 2.03). A single cysteine raises $S$ slightly above wild-type, whereas two or more cysteines reduce it, and no control approaches the enrichment of the top 10 ranking variants.
